# Zoogeochemical engineers: Galápagos giant tortoises link terrestrial and freshwater ecosystems through sediment transport and nutrient dynamics

**DOI:** 10.64898/2026.10.01.756004

**Authors:** Diego Ellis-Soto, Verónica Crespo-Pérez, Courtney L. Pike, Freddy Cabrera, Cristian Peñafiel, Herve Pierre-Paul, Christopher L. Dutton, David M. Post, Stephen Blake, Amanda L. Subalusky

## Abstract

Animal movements can transport materials across ecosystem boundaries, while simultaneously modifying habitats through physical engineering. We integrated GPS telemetry, camera trapping, sediment measurements and physicochemical sampling of ponds on Santa Cruz Island, Galápagos, to evaluate how giant tortoises connect and shape terrestrial and freshwater ecosystems. Telemetry revealed 2,254 visits by 21 tortoises to 12 ponds, with visits lasting up to 21.38 h and showing strong seasonal and diel variation, while camera traps showed peak pond use in January and during early morning and late afternoon. Greater tortoise abundance was associated with higher ammonium, dissolved organic carbon and total phosphorus concentrations, while ponds with persistently greater tortoise abundance had higher total suspended solids. Departing tortoises carried a mean 505.9 g of sediment, yielding a conservative potential export of 8.20 kg per tortoise per year and tens of tones annually. These findings identify giant tortoises as mobile zoogeochemical engineers linking terrestrial and freshwater ecosystems through physical and biogeochemical pathways.

## Introduction

Many ecosystems are connected through the spatial movement of organisms, material, and energy across ecosystem boundaries (Loreau et al., 2003). Linked meta-ecosystems can lead to emergent properties in stability, source-sink dynamics, and productivity that arise from these exchanges across donor and recipient ecosystems (Gounand et al., 2018; Leroux & Loreau, 2008; Loreau et al., 2003; Marleau et al., 2014; Polis et al., 1997). Animals in particular can exert strong impacts and shape ecosystems through their direct and indirect effects on elemental cycling and translocation, collectively described as zoogeochemistry (Earl & Zollner, 2017; Ellis-Soto et al., 2021; McInturf & others, 2019; Schmitz et al., 2018; Subalusky & Post, 2019; Vanni, 2002). Large-bodied or migratory animals in particular are able to cover large distances across elevational gradients and thus can connect distant ecosystems through elemental transport, enhancing their resilience and stability (Bauer & Hoye, 2014; Somveille & Ellis-Soto, 2022). However, animal migrations are disappearing worldwide, and large-bodied animals are particularly threatened by human activities, which reduce both their population numbers and their long distance movement abilities (Dirzo et al., 2014; Tucker et al., 2018; Wilcove & Wikelski, 2008). The role of these animals in maintaining ecological connections underscores the importance of understanding how their decline may influence ecosystem flows and connectivity amid global biodiversity loss.

Large animals can also shape ecosystems through physical engineering, independently of or in addition to transporting resources (Jones et al., 1994). Digging, trampling, wallowing and sediment disturbance can modify habitat structure, alter resource availability and create or maintain habitats used by other species (Meysman et al., 2006). These effects can be particularly important in freshwater systems, where physical modification by animals can influence water availability, sediment dynamics and habitat persistence (Moore, 2006; Naiman & Rogers, 1997). For example, feral donkeys in the Mojave and Sonoran deserts excavate groundwater-fed wells that increase surface-water availability, facilitate riparian tree establishment and increase vertebrate richness around these sites (Lundgren et al., 2021).

Some species simultaneously function as resource vectors and ecosystem engineers, potentially generating particularly strong effects on biogeochemical landscapes. For example, Pacific salmon (*Oncorhynchus sp.*) are one of the most well-known examples of animal resource vectors, annually transporting marine-derived nutrients in freshwater streams that can stimulate production (Naiman et al., 2002). However, they also dig redds on the stream benthos during reproduction, and this physical engineering disturbs algal biofilms and can decrease stream production, illustrating how both elemental transport and physical disturbance can interact (Holtgrieve & Schindler, 2011). This dual role is also illustrated by large herbivores in terrestrial– freshwater systems, where repeated movement across ecosystem boundaries can combine material transport with physical disturbance. Hippopotamus (*Hippopotamus amphibius*), for example, act as a conveyor belt between terrestrial grasslands and aquatic ecosystems, depositing large quantities of terrestrially derived organic matter and nutrients into rivers through excretion and egestion (Subalusky et al., 2015). These subsidies interact with system hydrology to strongly alter freshwater biogeochemistry and generate cascading effects on aquatic communities (Dutton et al., 2018). However, hippos also influence ecosystem hydrology by creating paths through riparian corridors and deepening water holes through their daily feeding migrations and wallowing (Naiman & Rogers, 1997). Together, these studies show that animals can alter freshwater ecosystems through the translocation of nutrients and organic matter, while also shaping the physical habitat through ecosystem engineering. Yet, the frequency and relative importance of these dual animal roles at the nexus of freshwater and terrestrial habitats has been comparatively understudied (Frauendorf et al., 2021) even as declining animal populations and movement patterns lead to their loss (Bauer & Hoye, 2014).

Early work on animal nutrient cycling and translocation primarily focused on fishes and birds (Kitchell et al., 1979), with more recent research on zoogeochemical processes increasingly focused on large mammals (Ferraro et al., 2026; Pringle et al., 2023; Schmitz et al., 2018). However, the ecological roles of reptiles such as turtles, tortoises, and crocodilians remain understudied (Lovich et al., 2018; Somaweera et al., 2020). Galápagos giant tortoises provide a compelling system for examining how a large terrestrial ectotherm can simultaneously transport resources and physically engineer habitats (Gibbs et al., 2010; Hunter et al., 2021). Galápagos giant tortoises are the largest terrestrial ectotherm worldwide and provide substantial ecosystem services through vegetation trampling, trail formation and soil compaction, seed dispersal and nutrient recycling (Blake et al., 2012; Ellis-Soto, 2020; Ellis-Soto et al., 2017; Gibbs et al., 2010).

With body masses exceeding 300 kg and historical densities reaching ∼2.5 individuals/ha, giant tortoises have the potential to influence ecosystem processes at broad spatial scales (Gibbs & Goldspiel, 2021). On Santa Cruz Island in the Galápagos, *Chelonoidis porteri* and *C. donfaustoi* form partially migratory populations, with altitudinal movements occurring primarily among adults and larger males showing the highest propensity to migrate (Bastille-Rousseau et al., 2019; Blake et al., 2013; Blake, Yackulic, et al., 2021; Yackulic et al., 2017). These movements can redistribute resources across elevational gradients.

Freshwater habitats, which occur primarily in the highlands, represent an important but poorly quantified component of these movements and their associated ecosystem effects. Giant tortoises repeatedly use ponds during their time in the highlands, potentially for drinking, wallowing and thermoregulation, creating multiple potential pathways of terrestrial–freshwater exchange. They may import terrestrially derived nutrients, organic matter and seeds through excretion and egestion, while exporting sediment that adheres to their carapaces during wallowing. Movements among ponds may also disperse aquatic plants, algae and macroinvertebrates (Ellis-Soto, 2020). Repeated entry, wallowing and departure may further disturb sediments, alter pond margins and help maintain open-water habitat (Froyd et al., 2014). Together, these processes suggest that freshwater ponds may function as nodes where tortoise movement generates both biogeochemical exchange and physical ecosystem engineering. As these ponds are the only freshwater ecosystems on the islands, these interactions may be particularly important in maintaining freshwater habitats and resources.

On Santa Cruz Island, these interactions occur across both protected and human-modified landscapes. Within the Galápagos National Park, many ponds are ephemeral and dry up during the dry season, whereas ponds in agricultural and touristic areas often receive sufficient rainfall to remain year-round, and others are created, maintained, and modified by people. Tortoises strongly select areas near ponds in agricultural landscapes (Pike et al., 2023), creating direct interactions among tortoise movement, freshwater availability and human land management. Human modification of pond occurrence and persistence may therefore alter not only where tortoises access freshwater, but also where their nutrient transport and ecosystem-engineering effects occur across the landscape. Palaeoecological evidence indicates that declines in giant tortoise populations on Santa Cruz coincided with transitions from open water to *Sphagnum* dominated wetlands, suggesting that tortoises may historically have helped maintain freshwater habitat structure (Froyd et al., 2014). Despite evidence that tortoises transport resources across landscapes, their ecological role in freshwater ecosystems remains poorly quantified, particularly their pond use and influence on freshwater biogeochemistry.

Here, we quantify how Galápagos giant tortoises connect terrestrial and freshwater ecosystems using multi-year GPS telemetry, camera trap monitoring, repeated measurements of pond physicochemical conditions and direct measurements of sediment transport. Specifically, we ask: (1) how tortoise pond use and revisitation vary across daily and seasonal timescales; (2) whether individual tortoises repeatedly use and connect multiple freshwater habitats; (3) whether tortoise abundance is associated with pond physicochemical and biogeochemical conditions; and (4) how much sediment tortoises export from ponds at individual and population scales.

## Material and methods

### Study site

The Galápagos climate is strongly seasonal, with a hot-wet season typically from January–May and a cool-dry season from June–December (Colinvaux, 1984). Freshwater ecosystems are scarce in Galápagos, and ponds are often the only source of water for wildlife, including giant tortoises, terrestrial birds, seabirds and livestock (Ellis-Soto pers. obs.). This study was conducted on Santa Cruz Island the most inhabited island in the archipelago, where ecosystems are heavily modified with non-native plants outnumbering native species, particularly at higher elevations (Guézou et al., 2010).

We sampled managed and natural ponds along the migratory routes of two giant tortoise species (*Chelonoidis porteri* and *C. donfaustoi*) at sites locally known as La Reserva and Cerro Fatal. Managed ponds occurred on private agricultural or touristic properties and were maintained or modified by people, whereas natural ponds occurred within the Galápagos National Park. Ponds were identified during long-term tortoise monitoring and fieldwork. A total of 21 ponds were studied across elevations ranging from 138-424 m, across the Galápagos National Park and agricultural areas from 2018-2025 (Figure 1, Suppl. Table 1), and one focal pond at Montemar was monitored with a time-lapse camera.

**Figure 1:**
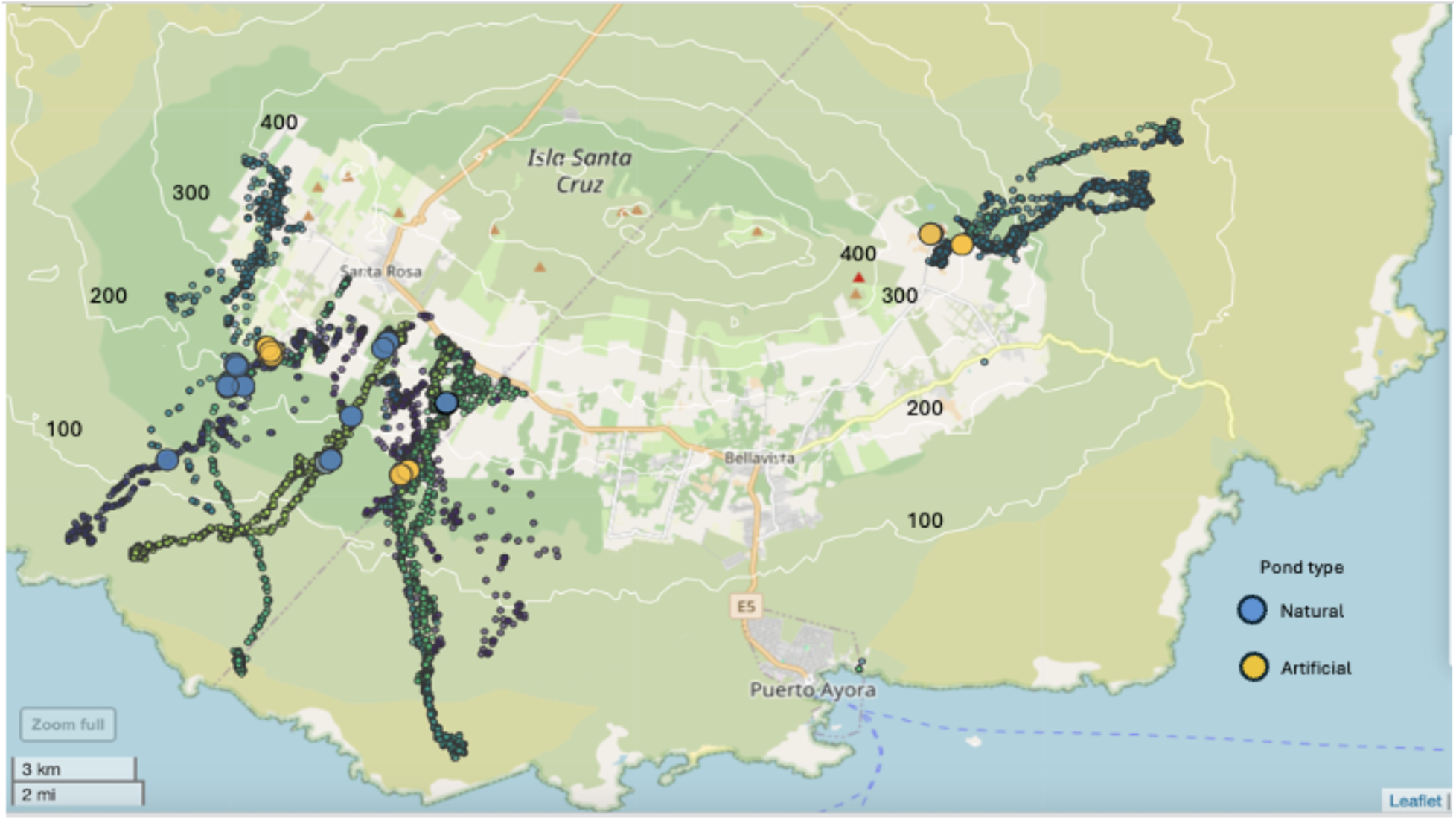
Migration routes of tracked *C. porteri* and *C. donfaustoi* individuals and pond locations on Santa Cruz Island. Sampling locations of ponds in Santa Cruz Island in the La Reserva and Cerro Fatal sites. Blue color ponds indicate natural ponds, while yellow circles represent managed ponds.

### Giant tortoise movement ecology and pond utilization

We analyzed tortoise pond use leveraging telemetry data from 14 individuals of *C. porteri* and seven individuals of *C. donfaustoi* from a long-term study of giant tortoise movement ecology (Bastille-Rousseau et al., 2019; Blake et al., 2013, 2024; Blake, Tapia, et al., 2021). Data were queried from the Movebank database (Kays et al., 2022) using the *move* package (Kranstauber et al., 2026) in R version 4.4.1. (R Core Team, 2024). Handling, tagging and GPS sampling rates are described in Blake et al. (2013).

To quantify daily and seasonal patterns of pond use, we identified visits by intersecting tortoise GPS locations with pond boundaries. Because pond extent varies over time, we approximated boundaries using GPS locations recorded during water-sampling visits. For 18 of the 21 ponds, we delineated pond extent as the area within 5 m of any sampling location. Three large ponds (Cerro Mesa, Laguna Chato, and Manzanillo 1) were digitized from high-resolution imagery and buffered by 5 m. Pond visits were identified using the *recurse* package in R (Bracis et al., 2018). Visits lasting ≤0.1 h were excluded to reduce brief pass-by events, and GPS locations falling within overlapping pond boundaries were counted only once. This yielded 2,254 unique pond visits by 21 tortoises at 12 ponds, which we used to summarize visit timing and duration. We constructed a weighted bipartite network linking individual tortoises to ponds, with edge weights representing the number of unique days each individual visited a given pond. To assess potential connectivity among freshwater habitats, we then identified successive detections of the same individual at different ponds and calculated the elapsed time between detections.

We quantified diel and seasonal pond use by tortoises using a time-lapse camera (Apeman model H40, Apeman International Company Ltd., Baldwin Park, CA) at Montemar Pond, located within an eco-resort at approximately 270 m elevation. The camera recorded one image every 15 min, and we manually counted tortoises observed in the water. Monitoring ran from 7 April 2022 to 11 January 2023, with a break from 20 August to 15 November 2022, yielding 18,235 observations.

### Physicochemical characteristics of ponds and nutrient sampling

Ponds were georeferenced using a Garmin i67 GPS and sampled from 2018-2025, yielding 157 sampling events across 21 ponds (Suppl. Table 1). At each visit, we counted the number of tortoises present and measured water temperature, conductivity, pH, and dissolved oxygen (DO) using a YSI 556 multiprobe (YSI Inc., Yellow Springs, Ohio). At each pond, we collected water samples for carbon and nutrient analysis. Water samples for dissolved inorganic nutrients (ammonium, NH_4_^+^-N; nitrate, NO_3_^-^-N and orthophosphate, PO_4_^-^-P) were filtered through a 0.2 µm Supor filter and frozen until analysis on a Seal AQ400 Discrete Analyzer at the University of Florida using standard protocols (Lipps et al., 2023). Water samples collected for total nitrogen (TN) and total phosphorus (TP) were unfiltered and frozen until analysis. They were digested using a potassium persulfate digestion and analyzed on a Seal AQ400 using standard protocols (Subalusky et al., 2018). Water samples for dissolved organic carbon (DOC) were filtered through a Whatman GF/C filter and frozen until analysis on a Shimadzu TOC analyzer for non-purgeable organic carbon (NPOC). The volume of water filtered was recorded, and filters were frozen for analysis of Chlorophyll. Chl *a* was analyzed using a basic methanol extraction, and extracts were analyzed on a Turner Trilogy fluorometer (Holm-Hansen & Riemann, 1978). We measured total suspended solids (TSS) by filtering a known volume of sample through a pre-weighed, pre-combusted Whatman GF/F filter, drying the filter at 60°C and re-weighing it.

### Multivariate pond conditions and nutrient associations

To characterize multivariate variation in pond conditions, we performed a principal component analysis (PCA) of water temperature, conductivity, dissolved oxygen, TN, TP, NPOC, ammonium and orthophosphate. Conductivity and nutrient concentrations were log(1 + x)-transformed, whereas temperature and dissolved oxygen remained untransformed; all variables were centred and scaled. The PCA included 67 complete sampling events from 21 ponds. Tortoise abundance (n = 66 events, 21 ponds), total suspended solids (n = 46, 19 ponds), chlorophyll-*a* (n = 37, 16 ponds) and elevation (n = 43, 19 ponds) were fitted independently as supplementary variables, so incomplete coverage did not reduce the ordination sample. All supplementary variables except elevation were log(1 + x)-transformed. Vector coordinates represent correlations with the first two PCA axes, tested using 4,999 permutations.

We tested associations between tortoise abundance and ammonium, nitrate, orthophosphate, TN, TP, NPOC, TSS and chlorophyll-a using separate Gamma generalized linear mixed-effects models with log links, each including tortoise abundance, pond type and season as fixed effects and pond identity as a random intercept. Tortoise abundance was quantified as the number of individuals observed at each pond during sampling and entered as log(1 + count), so that response ratios represent the proportional change in concentration per doubling of 1 + tortoise abundance. For each of our response variables, we additionally partitioned log-transformed tortoise abundance into between-pond and within-pond components, allowing us to distinguish temporal associations within ponds from persistent differences among ponds. Models were fitted using glmmTMB, and model adequacy was evaluated using simulation-based residual tests of uniformity and dispersion implemented in the DHARMa package in R (Brooks et al., 2017; Hartig, 2026). Full model estimates for all responses are shown in Suppl. Table 2.

### Physical ecosystem engineering of ponds

During each pond visit from 2018–2020, sediment was collected from the anterior portion of the tortoise carapace using a plastic spoon; from 2021 onwards, sediment was collected from both the anterior and posterior portions. Pre-2021 sediment quantities were therefore doubled to estimate whole-carapace loads. Tortoises were sampled opportunistically, with a maximum of one individual per pond visit, selecting the nearest tortoise in shallow water or observed exiting the pond to confirm recent pond use.

Samples were kept in a drying cabinet for up to one week until fully dry and weighed using a digital scale with an accuracy of 0.1 g. We obtained sediment-load measurements from 42 tortoises and carapace lengths from 26 individuals. We tested whether sediment loads differed between managed and natural ponds using a Tweedie generalized linear mixed-effects model with a log link and pond identity as a random intercept. We also tested the association between tortoise carapace length and dry sediment load using Pearson’s correlation.

To estimate potential annual sediment export, we integrated sediment loads with telemetry derived pond-departure rates. We included pond visits lasting more than 0.1 h and tortoises that had been tracked for at least 150 days. To avoid counting repeated exits and re-entries by the same tortoise on the same day as separate transport events, we counted a maximum of one departure per tortoise, pond and calendar day, for a total of 18,684 days across 20 tagged individual tortoises (51.2 tortoise-years). Annual departure rate was calculated as the number of departure events divided by total tracking effort.

Potential annual sediment export per tortoise was then estimated by multiplying this departure rate by the mean sediment load. We estimated uncertainty using 5,000 bootstrap iterations. In each iteration, tracked tortoises were resampled while retaining their departure counts and tracking effort, and sediment load measurements were resampled separately. Annual sediment export was recalculated in each iteration. We then scaled the individual-tortoise estimate to published estimates of mature population size (3,400 *Chelonoidis porteri* and 250 *C. donfaustoi*) (Cayot et al., 2017a,b). We scaled to mature individuals rather than to total population estimates to exclude juveniles and subadults that are unlikely to use ponds as these are at higher elevations than their range. Population sizes were fixed, and thus the confidence intervals reflect only variation among tracked tortoise revisitation rates and sediment load measurements.

## Results

### Giant tortoise seasonal and daily pond utilization

Our revisitation analysis identified 2,254 unique pond visits lasting >0.1 hours from 2018–2025 by 21 tortoises across 12 ponds. Median residence duration was 0.90 h (IQR = 1.01 h), with visits ranging from 0.10 to 21.38 h. Pond use was generally concentrated at individual ponds, although six tortoises visited more than one pond (Fig. 3), generating 47 observed transitions between ponds. Intervals between detections from one pond to a different pond ranged from 1.97 h to 0.88 years. A total of 182 visits (8.07% of all visits) extended through midnight, indicating that overnight pond use was relatively common (Fig. S1). Across tracked individuals, pond visitation was highest during the morning and evening and lowest around midday, peaking at 06:00 and reaching a minimum at 13:00 (Fig. 2). Camera monitoring at Montemar Pond showed a similar diel pattern, with greater tortoise abundance in the late afternoon and early morning and lower abundance around midday, consistent with frequent nighttime pond use (Fig. 4).

**Figure 2:**
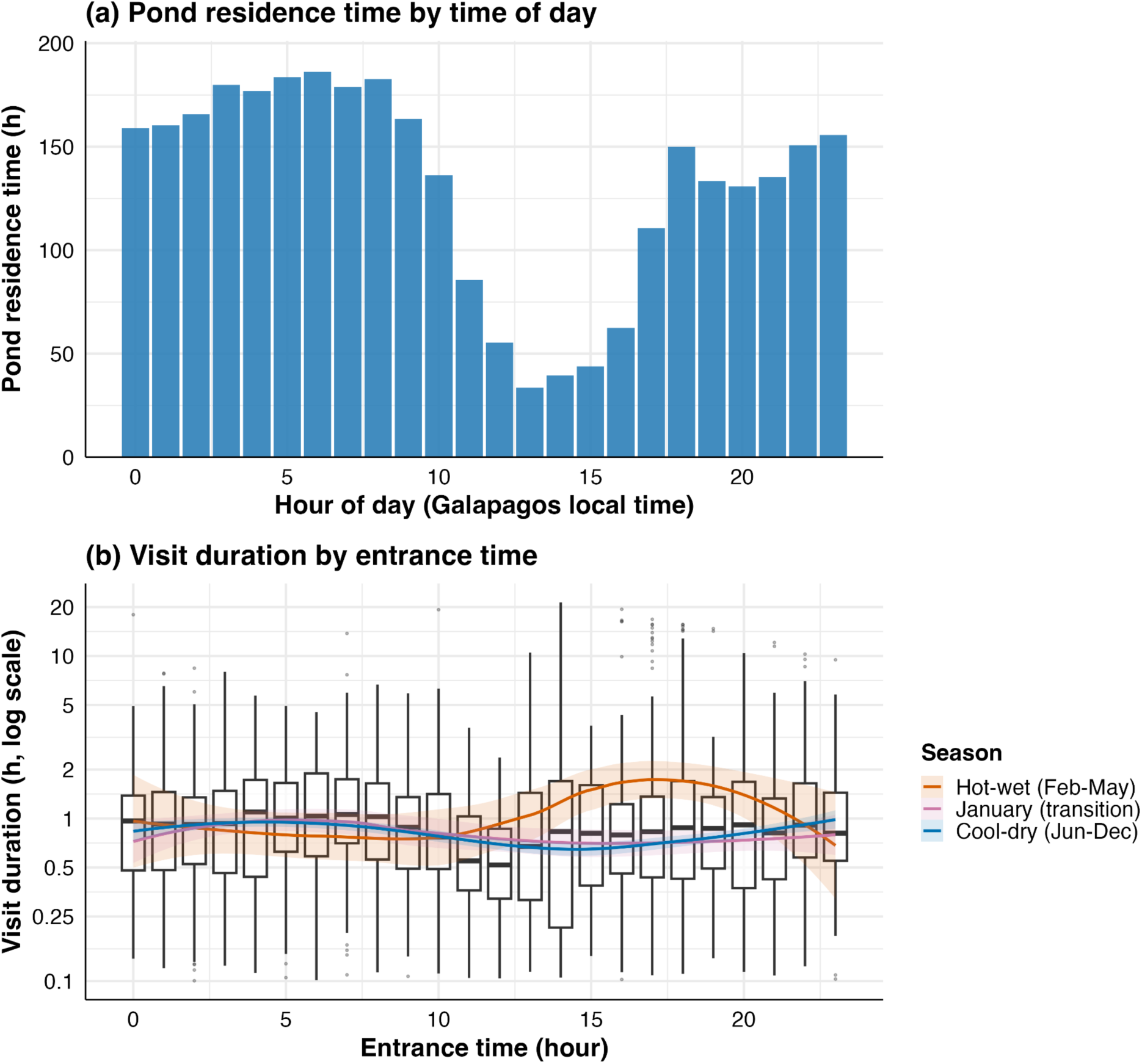
Movement-derived revisitation patterns of 21 tracked giant tortoises in freshwater ponds across Santa Cruz. (a) Total pond residence time by hour of day, summed across all visits. (b) Pond residence time by time of day.

**Figure 3:**
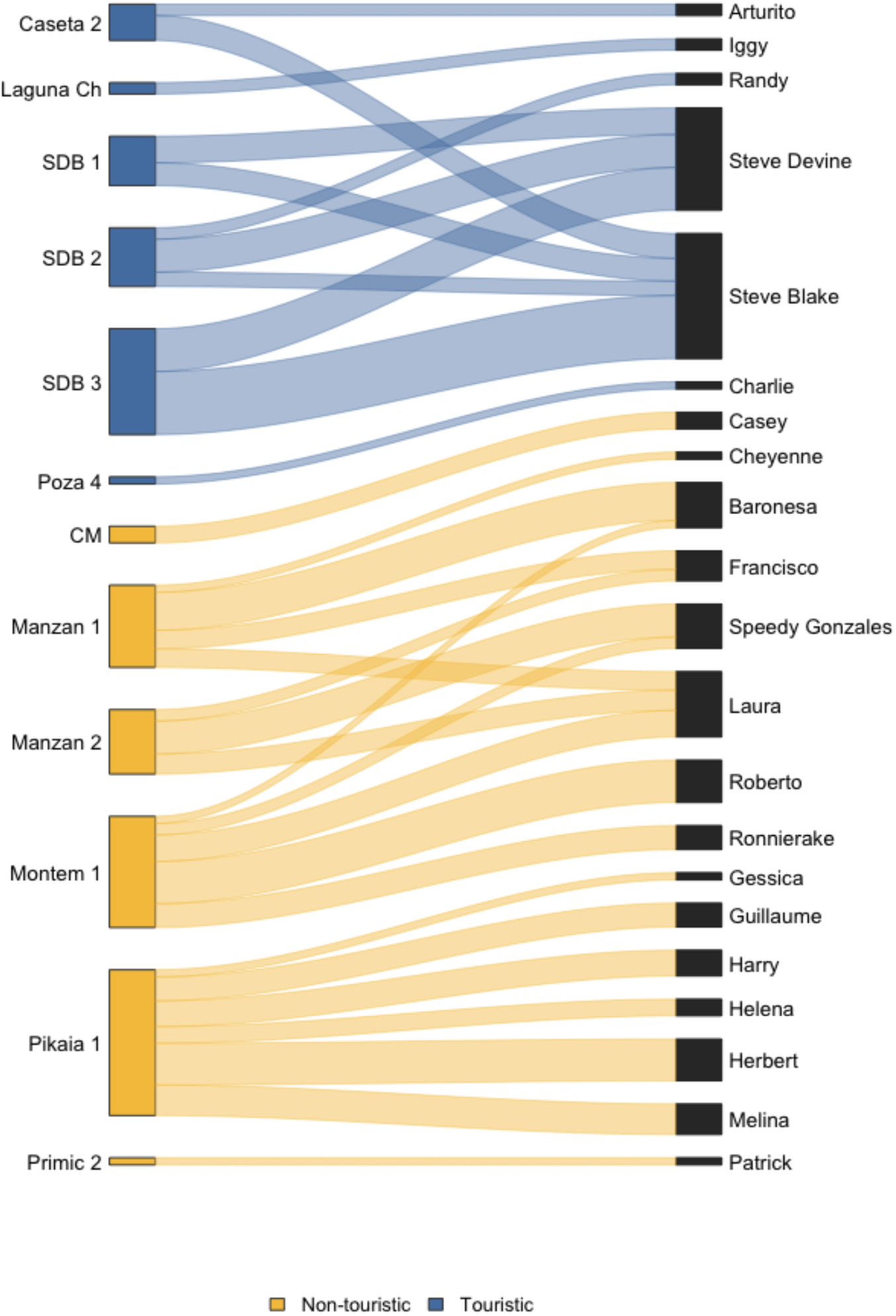
Recurse-based bipartite network of pond use by GPS-tracked Galápagos giant tortoises. Ponds are shown on the left and individual tortoises on the right. Ribbons indicate valid visit to a pond, and ribbon width is proportional to log(1 + the overall number of days a tortoise used a given pond). Pond nodes and their associated ribbons are coloured by pond category: blue indicates natural ponds and orange indicates managed ponds.

**Figure 4:**
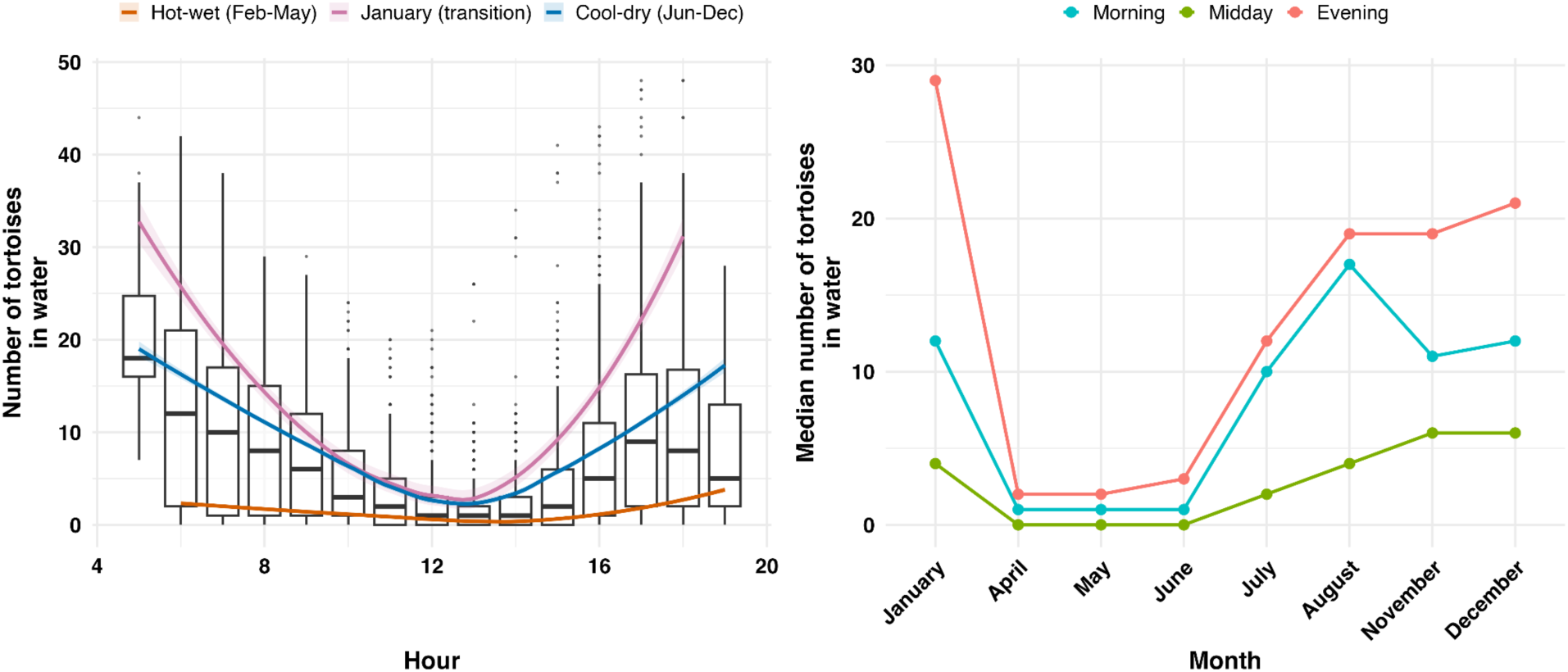
Daily and seasonal presence of tortoises at Montemar pond identified through camera monitoring. Giant tortoises show marked daily and seasonal revisitation patterns with highest numbers in the evening and morning, and lowest tortoise pond visitations during midday. (a) Number of tortoises in the water by hour of day. Boxplots give the median and interquartile range across all images taken in that hour; curves are loess fits with 95% confidence bands for each climatic season. (b) Median number of tortoises in the water by season and time of day, where morning is 05:00–11:59, midday 12:00–16:59 and evening 17:00–19:59. The Galápagos year divides into a hot-wet and a cool-dry season. January is shown separately as the transitional month, which carried the highest pond use. Monitoring ran from 7 April 2022 to 11 January 2023 with a break from 20 August to 15 November 2022.

Both camera and telemetry datasets indicated that pond use also varied seasonally. Telemetry showed that pond visitation increased from a minimum in April to a maximum in December. Camera trap monitoring at Montemar showed a broadly similar seasonal pattern, with mean pond abundance varying more than an order of magnitude among sampled months, from 1 tortoise in April to 13 in January. The magnitude of diel variation also differed between these months. From midday (12:00) to late afternoon (17:00), cameras showed mean abundance increased from 0.39 to 1.38 tortoises in April, an approximately 3.6-fold increase, compared to 3.48 to 24.8 tortoises in January, an approximately 7-fold increase.

### Pond environmental gradients and nutrient associations

Our PCA included 67 sampling events across 21 ponds. The first two axes explained 55.0% of the variation in pond water biogeochemistry (PC1 = 34.7%; PC2 = 20.3%; Fig. 6a). PC1 represented a gradient from lower to higher nutrient concentrations and conductivity, with nutrient-rich conditions occurring toward negative PC1. TN, TP and NPOC had the strongest loadings along this axis, with conductivity and ammonium also contributing. PC2 contrasted higher conductivity and ammonium with higher orthophosphate and total phosphorus. Among supplementary variables, total suspended solids was most strongly associated with the nutrient-rich, negative end of PC1 (PC1 correlation = −0.677; r^2^ = 0.485, P = 0.0002), followed by chlorophyll-a (r = −0.450; r^2^ = 0.207, P = 0.019) and tortoise abundance (r = −0.401; r^2^ = 0.171, P = 0.0020), whereas elevation showed no detectable association with the first two axes (r^2^ = 0.093, P = 0.135).

Mixed-effects models revealed positive associations between tortoise abundance and nutrient concentrations (Fig. 6b–d, Suppl. Table 2). After accounting for pond type, season and repeated sampling within ponds, each doubling of 1 + tortoise abundance was associated with 44% higher ammonium (response ratio = 1.44, 95% CI = 1.16–1.80, P < 0.001), with both within-pond (P = 0.012) and between-pond effects (P = 0.046) (Suppl. Fig. 2). NPOC was also positively associated with tortoise abundance (response ratio = 1.11, 95% CI = 1.02–1.20, P = 0.011), driven primarily by between-pond differences (P = 0.005). Total phosphorus showed a weaker but still significant association with tortoise abundance (response ratio = 1.19, 95% CI = 1.03–1.37, P = 0.021), whereas TN was marginal and nitrate and orthophosphate showed no detectable associations (Suppl. Table 2). Chlorophyll-a was not associated with tortoise abundance (response ratio = 0.92, 95% CI = 0.73–1.15, P = 0.452), indicating no immediate algal response to nutrient enrichment. Total suspended solids showed a different pattern: event-level analysis showed no significant association with tortoise abundance (response ratio = 1.15, 95% CI = 0.92–1.42, P = 0.214), but between-pond partitioning revealed significant effects (response ratio = 1.45, 95% CI = 1.08–1.96, P = 0.013). TSS was 3.42-fold higher in managed than natural ponds (95% CI = 1.37–8.55, P = 0.009), potentially reflecting substrate differences and human management rather than tortoise activity.

### Physical ecosystem engineering of ponds

The median weight of dry sediment transported out of ponds by a single tortoise was 340 g (μ=505.9, SD = 518.9, range of 0-2,112 g, Fig. 5). This estimate is likely conservative, as we were not able to collect all sediment from the tortoises’ bodies. Sediment loads were 2.02 times higher at managed ponds than natural ponds inside the national park (marginally significant P = 0.050, 95% CI = 1.00–4.10). Carapace length was not significantly associated with dry sediment load (n = 26; Fig. S3). Across 51.2 years of accumulated tortoise tracking data, we recorded 829 unique tortoise–pond departure days equivalent to ∼16 departures per tortoise per year. Combining this rate with the distribution of our sediment load data yielded a conservative estimate of 8.20 kg of dry sediment transported out of ponds per tortoise per year (95% bootstrap CI = 2.42–18.08 kg). When scaled for total adults in current populations, this yielded approximately 27.9 tones per year for C. porteri (95% bootstrap CI = 8.2–61.5 tones yr⁻¹) and 2.05 tones per year for C. donfaustoi (95% CI = 0.61–4.52 tones yr⁻¹)(Suppl. Fig. 4).

**Figure 5:**
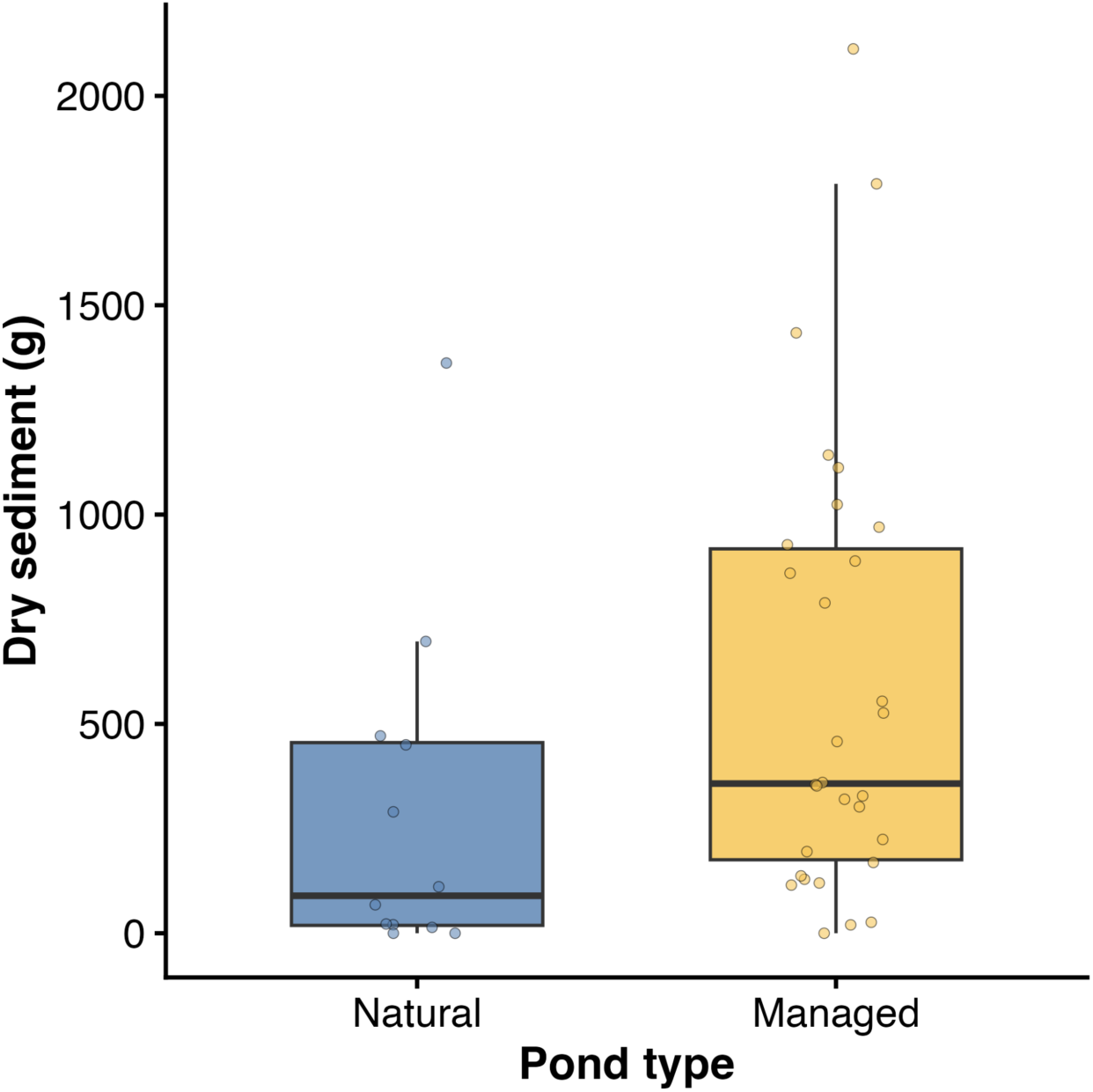
Sediment transport of Galápagos giant tortoises out of freshwater ponds. Larger quantities of sediment were obtained from tortoise carapaces in managed ponds when compared to natural ponds in the Galápagos National Park.

**Figure 6:**
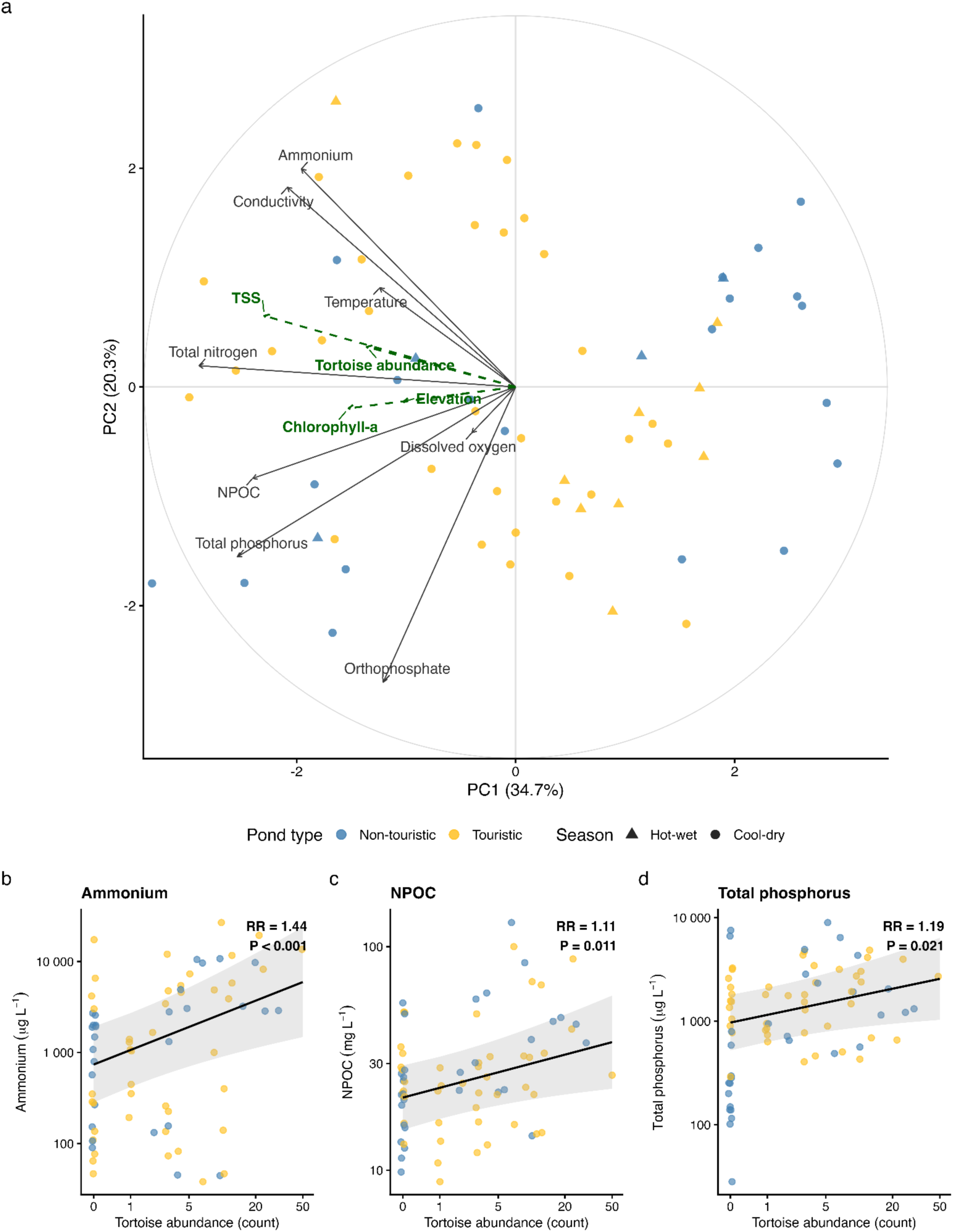
Pond physicochemical conditions and their associations with tortoise abundance. (a) Principal component analysis (PCA) of 67 sampling events from 21 ponds. Points represent sampling events, coloured by pond type and shaped by season. Solid vectors represent physicochemical variables used to construct the PCA, whereas dashed vectors represent supplementary variables fitted after ordination. Vector direction indicates increasing values and vector length indicates the strength of association with the first two PCA axes. (b–d) Model-adjusted relationships between contemporaneous tortoise abundance and (b) ammonium, (c) NPOC, (d) total phosphorus. Lines show predictions from Gamma mixed-effects models accounting for pond type, season and repeated sampling within ponds. Points represent individual pond-sampling events.

## Discussion

Galápagos giant tortoises repeatedly used freshwater ponds, with strong diel and seasonal variation, prolonged residence times and movement among multiple ponds by some individuals. Greater tortoise abundance was associated with nutrient-rich pond conditions, particularly higher ammonium concentrations, and higher TSS among ponds, whereas chlorophyll-*a* showed no detectable response. Departing tortoises also transported substantial quantities of sediment from ponds into surrounding terrestrial environments, potentially exporting several kilograms per tortoise per year and tens of tones per year at the species level. Together, these findings show that repeated tortoise movements across pond boundaries generate both biogeochemical and physical connectivity between terrestrial and freshwater ecosystems, with potential consequences for freshwater habitat structure and maintenance in Galápagos.

### Giant tortoise behavior, pond use and connectivity

Individual tortoise pond use was generally concentrated at a single pond or pond complex, although several individuals used multiple ponds, providing a mechanism for connecting otherwise discrete freshwater habitats. Some visits lasted more than 21 h, increasing opportunities to deposit nutrients and organic matter, disturb sediments and transport material between ponds and surrounding terrestrial habitats. Successive detections of the same tortoise at different ponds occurred within the same day, while repeated revisitation indicates that ponds are persistent features of individual movement routines.

Pond use was lowest around midday and greatest during the morning and late afternoon or evening, with both telemetry and camera data showing similar seasonal patterns. Telemetry-derived visitation increased through the latter half of the year and peaked in December, while camera observations at Montemar were highest in January and lowest in April. These patterns are consistent with seasonal migration in which tortoises move downslope into the lowlands, away from most ponds, during the hot-wet season in response to rainfall and resource availability (Bastille-Rousseau et al., 2019; Blake et al., 2013; Yackulic et al., 2017). Together, these diel and seasonal patterns suggest that nutrient deposition, wallowing and sediment disturbance are concentrated during cooler parts of the day and may occur as seasonal pulses rather than being evenly distributed throughout the year.

### Giant tortoise abundance, nutrient dynamics and primary producer responses

Greater tortoise abundance was associated with the nutrient rich end of the pond-chemistry gradient and with higher ammonium concentrations. Because tortoise abundance was included as a supplementary variable in the PCA, the ordination indicates association rather than habitat preference. Positive associations between tortoise abundance and pond nutrient conditions are consistent with animal-mediated nutrient inputs through excretion and egestion documented in other species (Ferraro et al., 2024, 2026; Subalusky et al., 2015); but may also arise through bioturbation and sediment disturbance. The association was strongest and significant for ammonium, with each doubling of 1 + tortoise abundance associated with 44% higher concentrations.

Total phosphorus and NPOC were also significantly associated with tortoise abundance. In contrast, total nitrogen was only marginally associated with tortoise abundance, and nitrate and orthophosphate showed no detectable associations (Suppl. Table 2). Total suspended solids followed a different pattern: Concentrations showed no detectable association to tortoise abundance at the level of individual sampling events, but ponds with persistently higher tortoise abundance had higher suspended sediment (between-pond response ratio = 1.45, P = 0.013), consistent with repeated wallowing and sediment disturbance maintaining elevated turbidity over time rather than producing detectable pulses at the moment of sampling. Suspended solids were also 3.42-fold higher in managed than natural ponds, so substrate and pond management may also contribute to these patterns. Together, these results suggest that greater tortoise abundance was associated with higher nutrient concentrations, particularly ammonium, likely through excretion and egestion or mineralization of animal-derived organic matter. However, wallowing and sediment disturbance could additionally release pore-water ammonium into the water column (Vanni, 2002).

Chlorophyll-a was not significantly associated with tortoise abundance, suggesting that nutrient enrichment did not translate into algal biomass increases. Primary producer responses may depend on light (potentially limited by the elevated TSS in managed ponds), hydrology, nutrient ratios, grazing, and temporal lags. The contrast between strong ammonium signatures and no significant chlorophyll response suggests an important decoupling. Tortoises could be actively subsidizing ponds with nutrients and organic matter, yet these biogeochemical changes may not immediately manifest as primary production, possibly due to decreased light availability due to tortoise bioturbation. Instead, tortoise-mediated carbon and nutrient inputs likely support alternative food-web pathways, including macroinvertebrate communities, particularly coprophagous larvae, while tortoise wallowing and sediment transport fundamentally reshape pond structure, margins, and sediment dynamics.

### Physical ecosystem engineering and sediment export

Our study identified a direct physical pathway through which giant tortoises may engineer freshwater habitats by transporting pond sediment into surrounding terrestrial environments. Tortoises carried a mean of 505.9 g of dry sediment during pond departures, corresponding to a conservative potential export of 8.20 kg per tortoise per year and tens of tones annually at the population scale. These estimates are likely conservative because the large body size of giant tortoises prevented complete sampling of sediment from the entire body, with measurements restricted to accessible portions of the carapace.

Our findings extend the established role of giant tortoises as ecosystem engineers through trampling, trail formation and vegetation modification to include freshwater sediment transport. Repeated entry, wallowing and departure may alter pond margins, access routes, sediment deposition and open-water persistence, consistent with palaeoecological evidence from Santa Cruz (Froyd et al., 2014). Similar coupled physical and biogeochemical effects occur in other megavertebrate systems: hippopotamus movements and wallowing connect terrestrial grasslands with rivers while altering aquatic biogeochemistry and habitat structure (Dutton et al., 2018; Subalusky et al., 2015), and African forest elephants modify trails, soil and water resources, and forest structure (Keany et al., 2024; Metsio Sienne et al., 2014). In our system, sediment loads and total suspended solids were higher in managed ponds than in natural ponds. Managed ponds often contain deeper accumulations of soil and mud, whereas many natural ponds have exposed lava bottoms (F. Cabrera, pers. obs.), suggesting that substrate and management may influence the magnitude and spatial distribution of tortoise engineering effects.

Our sediment estimates represent potential fluxes rather than measured ecosystem-wide budgets, and the broad bootstrap intervals reflect variation among tracked individuals and sediment loads. Camera monitoring was also concentrated at a single site, and the nutrient relationships remain observational. However, these estimates provide evidence that tortoise activity and movement could have substantial geomorphic impacts on freshwater ponds and warrant further study.

### Zoogeochemical engineering, restoration and management implications

Nutrient cycling remains a major gap in understanding the ecological effects of tortoises (Falcón & Hansen, 2018; Lovich et al., 2018), reflecting the broader emphasis of zoogeochemical research on mammals and birds. Studies from South America are also rare, accounting for only 4% of studies in a recent synthesis of mammalian zoogeochemistry. Our findings extend this body of research to a large reptile and demonstrate how animal movement can couple biogeochemical transport with physical ecosystem engineering (Ellis-Soto et al., 2021; Ferraro et al., 2024; Schmitz et al., 2018).

Giant tortoises can simultaneously transport nutrients, organic matter and propagules into ponds while exporting sediment to surrounding terrestrial environments, thereby linking multiple ecosystem processes across terrestrial–freshwater boundaries. Our telemetry also shows that some individuals revisit multiple ponds across seasons, potentially dispersing aquatic plants, invertebrates and seeds among otherwise isolated ponds (Ellis-Soto, 2020). More broadly, biologging can reveal how individual movement connects habitats and environmental conditions in ways relevant to conservation and management (Ellis-Soto et al. 2025). Together, these pathways identify tortoises as mobile zoogeochemical engineers with implications for restoration and rewilding. Historical declines in giant tortoises likely reduced freshwater-associated processes such as sediment disturbance and material redistribution, whereas population recovery and reintroduction may restore some of these functions (Falcón & Hansen, 2018; Froyd et al., 2014; Tapia Aguilera & Gibbs, 2023). Future work should quantify how wallowing and sediment removal alter pond morphology and persistence, estimate nutrient inputs through excretion and egestion, and experimentally test how tortoise abundance influences freshwater nutrient dynamics. Comparisons across natural and artificial ponds could further determine how human water provisioning modifies these effects. Because these habitats support waterbirds, aquatic invertebrates, rare plants and other freshwater-associated organisms (Ellis-Soto, 2020), tortoise-mediated changes to pond structure and persistence may have broader consequences for freshwater biodiversity. Resolving these mechanisms will become increasingly important as tortoise populations recover or are reintroduced across Galápagos, where decisions about pond design, water provisioning and surrounding vegetation may shape where their ecosystem-engineering effects are expressed.

## Supplementary Material

**Supplementary Figure 1:**
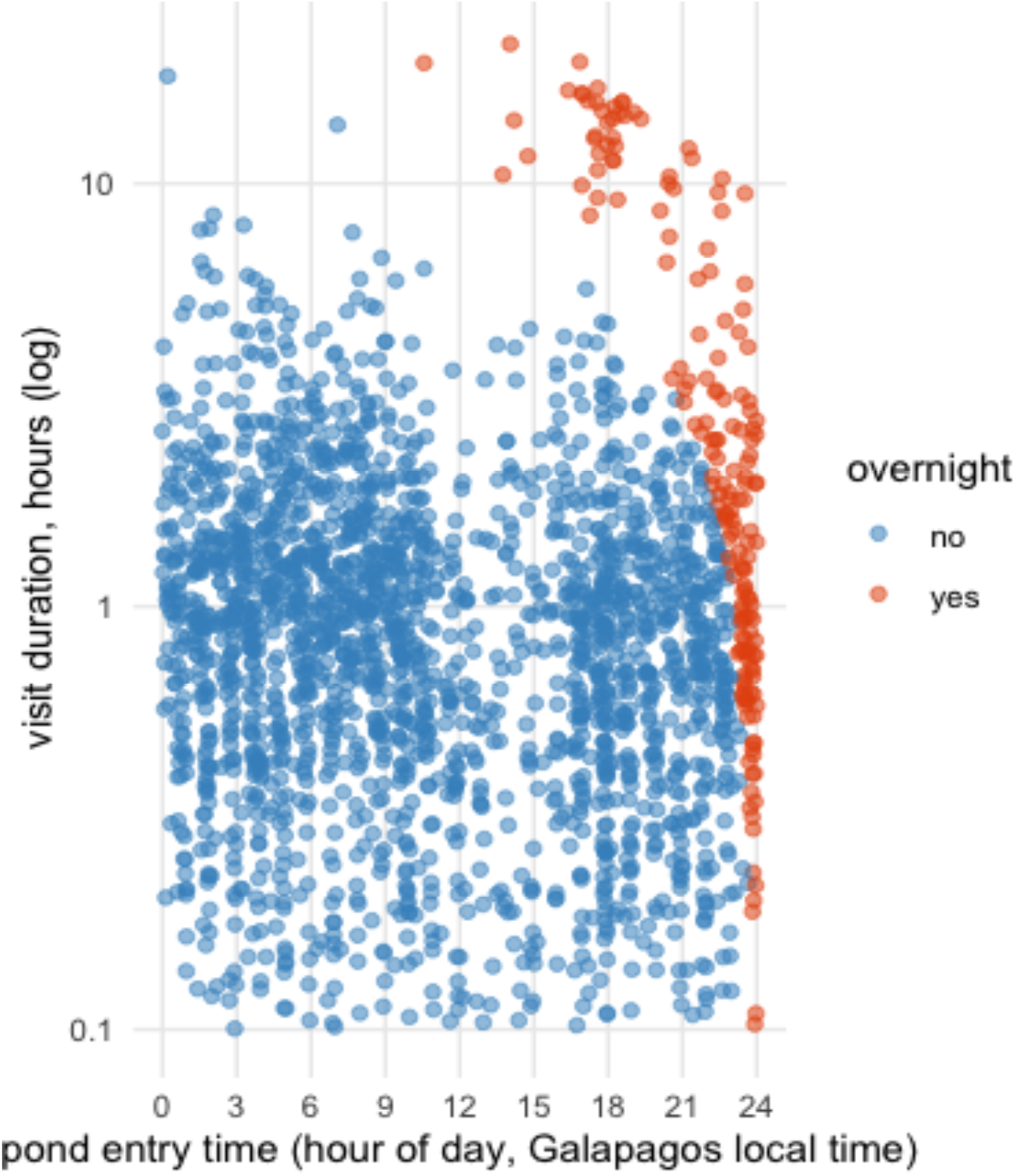
Evidence of overnight pond use by Galápagos giant tortoises. Relationship between pond-entry time and visit duration, with visits classified according to whether they extended past midnight. Each point represents an individual pond visit; colours distinguish overnight from non-overnight visits.

**Supplementary Figure 2:**
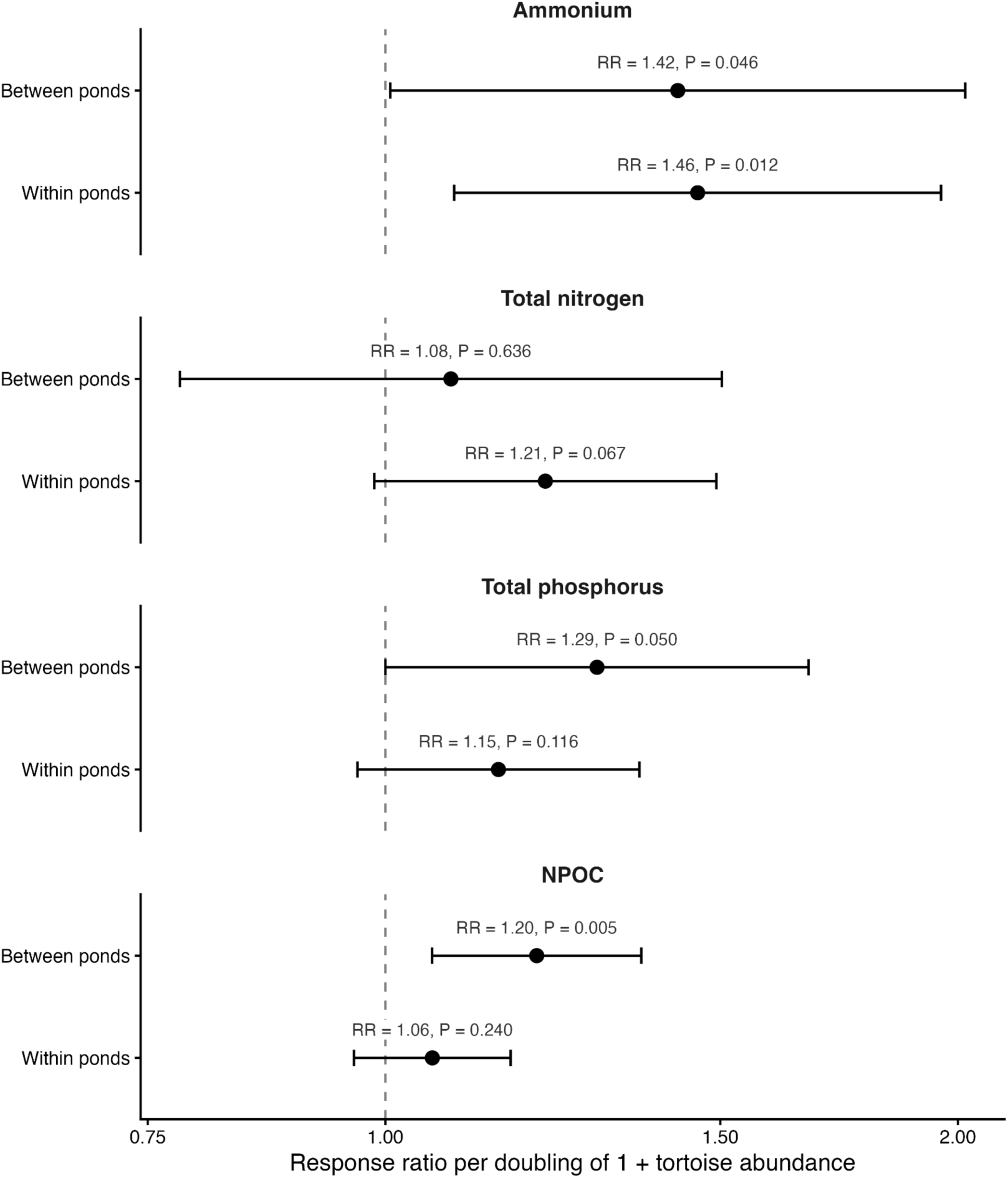
Within- and between-pond associations between tortoise abundance and pond nutrient concentrations. Tortoise abundance was partitioned into a between-pond component (the mean for each pond) and a within-pond component (the deviation of each sampling event from that pond mean), so that persistent differences among ponds could be distinguished from temporal variation within them. Points show response ratios for a doubling of 1 + tortoise abundance, estimated from Gamma generalized linear mixed-effects models accounting for pond type, season and repeated sampling within ponds; bars show 95% confidence intervals and the dashed line at 1 indicates no association.

**Supplementary Figure 3:**
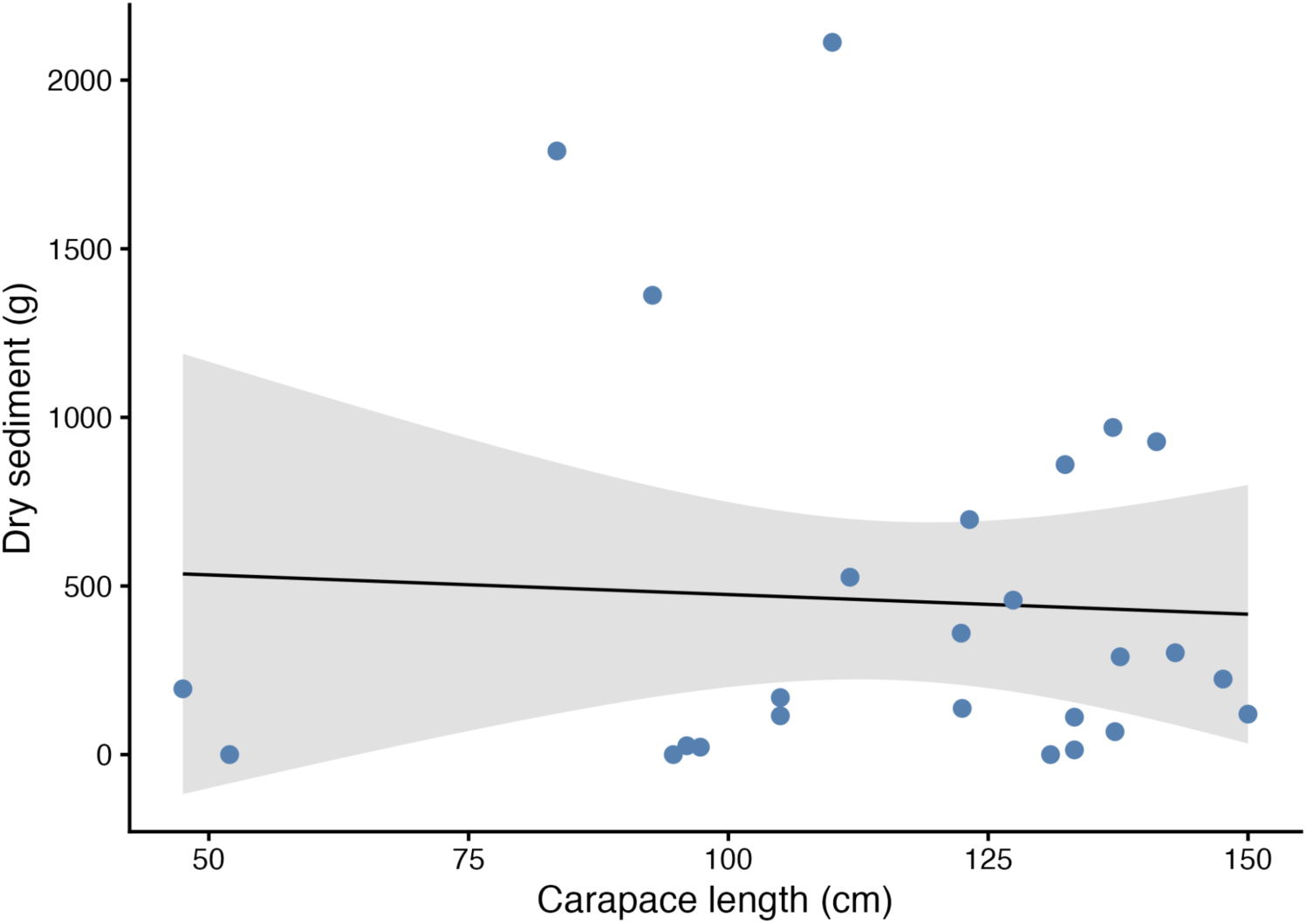
Relationship between tortoise carapace length and dry sediment mass in grams scraped from each individual tortoise.

**Supplementary Figure 4:**
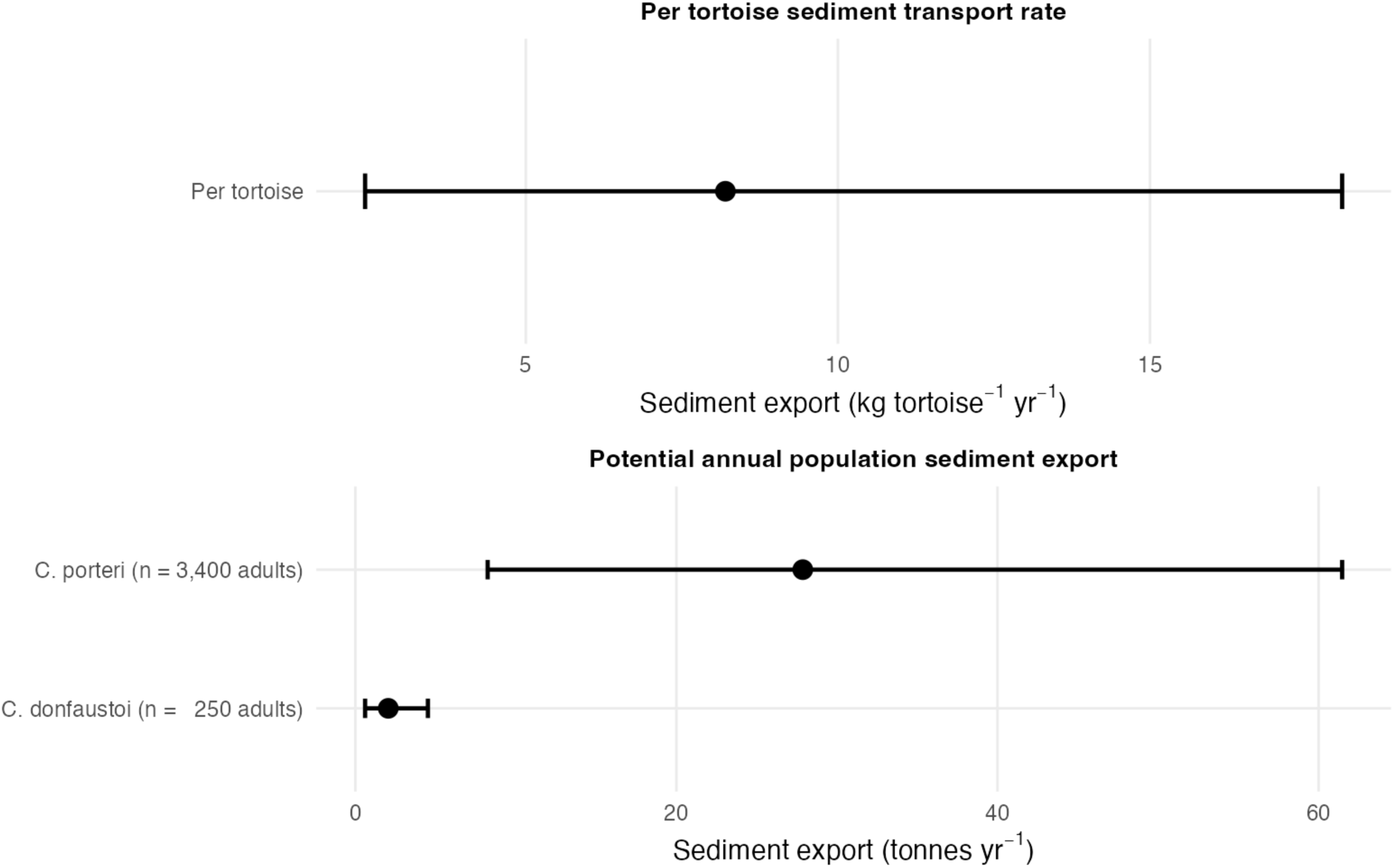
Potential sediment export by Galápagos giant tortoises. (a) Per-tortoise export rate (8.20 kg yr⁻¹, 95% CI 2.42–18.08). (b) Export scaled to population estimate as an adults-only scenario (Chelonoidis porteri, n = 3,400 adults; C. donfaustoi, n = 250 adults). Points show estimates and bars 95% bootstrap confidence intervals.

**Supplementary Table 1:**
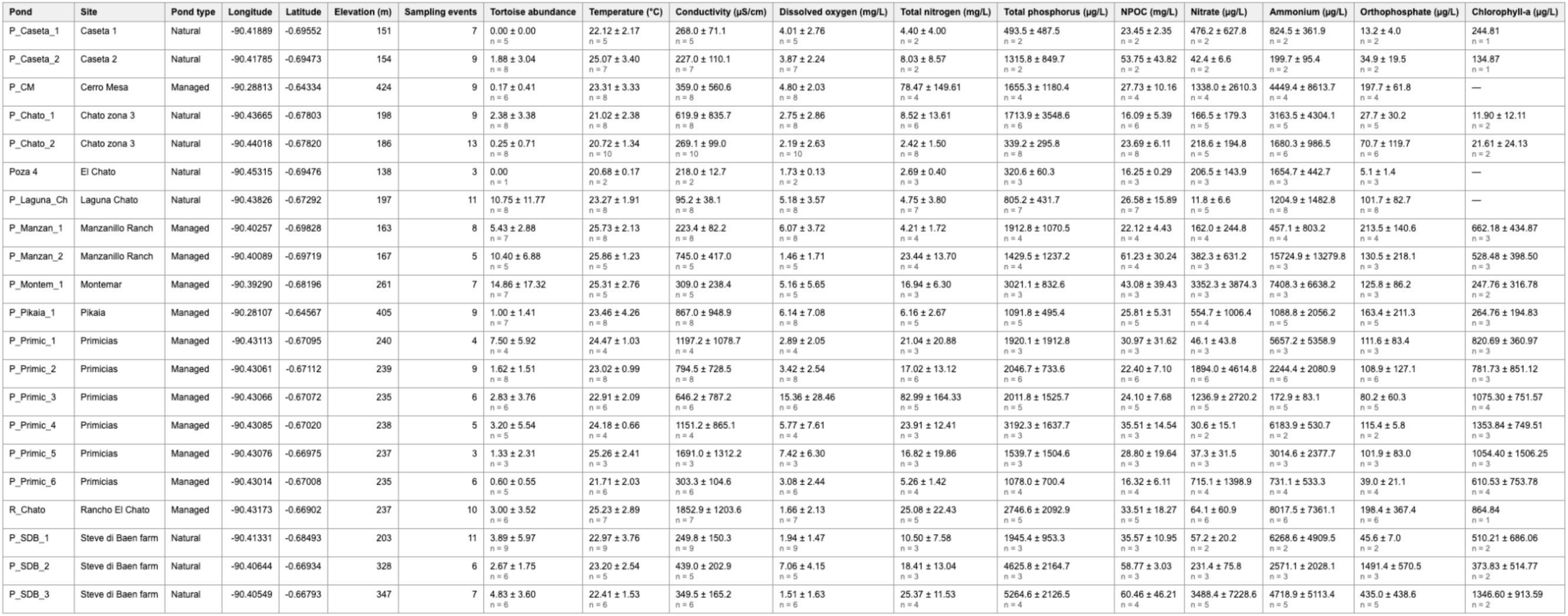
Sampling effort, tortoise abundance and physicochemical characteristics of the 21 freshwater ponds studied on Santa Cruz Island, 2018–2025. Elevation is taken from the SRTM digital elevation model at each pond’s coordinates. Tortoise abundance and water chemistry are given as mean ± SD (n), where n is the number of sampling events with that measurement. Dashes mark variables never measured at that pond

**Supplementary Table 2:** Associations between tortoise abundance and pond water chemistry. Response ratios give the proportional change in concentration per doubling of 1 + tortoise abundance, estimated from Gamma generalized linear mixed-effects models with pond type and season as fixed effects and pond identity as a random intercept. Between- and within-pond columns partition tortoise abundance into each pond’s mean and the deviation of each sampling event from that mean.

| Response | Sampling events | Ponds | Response ratio (95% CI) | P | Between ponds (95% CI) | P | Within ponds (95% CI) | P |
| --- | --- | --- | --- | --- | --- | --- | --- | --- |
| Ammonium | 73 | 21 | 1.44 (1.16-1.80) | <0.001 | 1.42 (1.01-2.02) | 0.046 | 1.46 (1.09-1.96) | 0.012 |
| Total nitrogen | 76 | 21 | 1.18 (0.98-1.41) | 0.080 | 1.08 (0.78-1.50) | 0.636 | 1.21 (0.99-1.49) | 0.067 |
| Nitrate | 69 | 21 | 1.05 (0.72-1.54) | 0.786 | 1.00 (0.59-1.70) | 0.995 | 1.11 (0.66-1.87) | 0.693 |
| Total phosphorus | 76 | 21 | 1.19 (1.03-1.37) | 0.021 | 1.29 (1.00-1.67) | 0.050 | 1.15 (0.97-1.36) | 0.116 |
| Orthophosphate | 73 | 21 | 0.92 (0.74-1.15) | 0.466 | 1.29 (0.88-1.89) | 0.186 | 0.81 (0.63-1.04) | 0.097 |
| NPOC | 77 | 21 | 1.11 (1.02-1.20) | 0.011 | 1.20 (1.06-1.36) | 0.005 | 1.06 (0.96-1.16) | 0.240 |
| TSS | 53 | 20 | 1.15 (0.92-1.42) | 0.214 | 1.45 (1.08-1.96) | 0.013 | 0.94 (0.72-1.24) | 0.682 |
| Chlorophyll-a | 43 | 18 | 0.92 (0.73-1.15) | 0.452 | 0.97 (0.68-1.39) | 0.887 | 0.87 (0.63-1.20) | 0.396 |

## Acknowledgements

DES acknowledges support from the Rufford Foundation, the Yale Institute of Biospheric Studies, the Presidential Postdoctoral Fellowship Program and NASA FINESST (80NSSC22K1535). This study was framed under the Research Permit PC-35-18, PC-16-19, PC-28-20, PC-17-21, PC-25-22, PC-37-23, PC-25-24, and PC-41-25 issued by the Galápagos National Park. This publication is contribution number 2867 of the Charles Darwin Foundation for the Galápagos Islands. DMP, CLD, and ALS were supported by the National Science Foundation (DEB 1354053 and 175372). CLD and ALS were also supported by the University of Florida Biodiversity Institute. Funding through the Galápagos Tortoise Movement Ecology Program was provided by the Houston Zoo, Galápagos Conservation Trust, Saint Louis Zoo, the COmON Foundation, and a bequest to the Galápagos Tortoise Movement Ecology Program. Thank you to the Charles Darwin Foundation staff for assistance in administration and permitting; we are grateful to Ainoa Nieto, Jose Haro and Partick Moldowan for providing logistical support. We thank the landowners of Montemar, Primicias, El Chato and Manzanillo for access to their pond during this study. The authors declare no conflict of interest.

## References

Bastille-Rousseau, G., Yackulic, C. B., Gibbs, J. P., Frair, J. L., Cabrera, F., & Blake, S. (2019). Migration triggers in a large herbivore: Galápagos giant tortoises navigating resource gradients on volcanoes. Ecology, 100(6), e02658.

Bauer, S., & Hoye, B. J. (2014). Migratory animals couple biodiversity and ecosystem functioning worldwide. Science, 344(6179), 1242552.

Blake, S., Cabrera, F., Cruz, S., Ellis-Soto, D., Yackulic, C. B., Bastille-Rousseau, G., Wikelski, M., Kuemmeth, F., Gibbs, J. P., & Deem, S. L. (2024). Environmental variation structures reproduction and recruitment in long-lived mega-herbivores: Galapagos giant tortoises. Ecological Monographs, 94(2), e1599. 10.1002/ecm.1599

Blake, S., Tapia, P. I., Safi, K., & Ellis-Soto, D. (2021). Diet, behavior, and activity patterns. In Galapagos giant tortoises (pp. 207–239). Elsevier.

Blake, S., Wikelski, M., Cabrera, F., Guezou, A., Silva, M., Sadeghayobi, E., Yackulic, C. B., & Jaramillo, P. (2012). Seed dispersal by Galápagos tortoises. Journal of Biogeography, 39(11), 1961–1972.

Blake, S., Yackulic, C. B., Cabrera, F., Deem, S. L., Ellis-Soto, D., Gibbs, J. P., Kummeth, F., Wikelski, M., & Bastille-Rousseau, G. (2021). Movement ecology. In Galapagos giant tortoises (pp. 261–279). Elsevier.

Blake, S., Yackulic, C. B., Cabrera, F., Tapia, W., Gibbs, J. P., Kümmeth, F., & Wikelski, M. (2013). Vegetation dynamics drive segregation by body size in Galapagos tortoises migrating across altitudinal gradients. Journal of Animal Ecology, 82(2), 310–321.

Bracis, C., Bildstein, K. L., & Mueller, T. (2018). Revisitation analysis uncovers spatio-temporal patterns in animal movement data. Ecography, 41(11), 1801–1811.

Brooks, M. E., Kristensen, K., van Benthem, K. J., Magnusson, A., Berg, C. W., Nielsen, A., Skaug, H. J., Maechler, M., & Bolker, B. M. (2017). glmmTMB Balances Speed and Flexibility Among Packages for Zero-inflated Generalized Linear Mixed Modeling. The R Journal, 9(2), 378–400. 10.32614/RJ-2017-066

Cayot, L. J., Gibbs, J. P., Tapia, W., & Caccone, A. (2017a). Chelonoidis porteri. The IUCN Red List of Threatened Species 2017: e.T9026A82777132.

Cayot, L. J., Gibbs, J. P., Tapia, W., & Caccone, A. (2017b). Chelonoidis donfaustoi. The IUCN Red List of Threatened Species 2017: e.T90377132A90377135.

Colinvaux, P. A. (1984). The Galápagos climate: Present and past.

Dirzo, R., Young, H. S., Galetti, M., Ceballos, G., Isaac, N. J., & Collen, B. (2014). Defaunation in the Anthropocene. Science, 345(6195), 401–406.

Dutton, C. L., Subalusky, A. L., Hamilton, S. K., Rosi, E. J., & Post, D. M. (2018). Organic matter loading by hippopotami causes subsidy overload resulting in downstream hypoxia and fish kills. Nature Communications, 9, 1951.

Earl, J. E., & Zollner, P. A. (2017). Advancing research on animal-transported subsidies by integrating animal movement and ecosystem modelling. Journal of Animal Ecology, 86(5), 987– 997.

Ellis-Soto, D. (2020). Giant tortoises connecting terrestrial and freshwater ecosystems in Santa Cruz Island. Galapagos Giant Tortoises, 286.

Ellis-Soto, D., Blake, S., Soultan, A., Guézou, A., Cabrera, F., & Lötters, S. (2017). Plant species dispersed by Galapagos tortoises surf the wave of habitat suitability under anthropogenic climate change. PLoS One, 12(7), e0181333.

Ellis-Soto, D., Ferraro, K. M., Rizzuto, M., Briggs, E., Monk, J. D., & Schmitz, O. J. (2021). A methodological roadmap to quantify animal-vectored spatial ecosystem subsidies. Journal of Animal Ecology, 90(7), 1605–1622.

Ellis-Soto, D., Flack, A., Strandburg-Peshkin, A., Wild, T. A., Williams, H. J., & O’Mara, M. T. (2025). From biologging to conservation: Tracking individual performance in changing environments. Proceedings of the National Academy of Sciences, 122(31), e2410947122. 10.1073/pnas.2410947122

Falcón, W., & Hansen, D. M. (2018). Island rewilding with giant tortoises in an era of climate change. Philosophical Transactions of the Royal Society B: Biological Sciences, 373(1761), 20170442. 10.1098/rstb.2017.0442

Ferraro, K. M., Albrecht, D., Hendrix, J. G., Vander Wal, E., Schmitz, O. J., Webber, Q. M. R., & Bradford, M. A. (2024). The biogeochemical boomerang: Site fidelity creates nutritional hotspots that may promote recurrent calving site reuse. Ecology Letters, 27, e14491. 10.1111/ele.14491

Ferraro, K. M., Forbes, E. S., Abraham, A. J., & Monk, J. D. (2026). Hot spots or hot moments? Contextualizing the spatio-temporal scale of research on animal inputs. Ecography, 2026(6), e08351. 10.1002/ecog.08351

Frauendorf, T. C., Subalusky, A. L., Dutton, C. L., Hamilton, S. K., Masese, F. O., Rosi, E. J., Singer, G. A., & Post, D. M. (2021). Animal legacies lost and found in river ecosystems. Environmental Research Letters, 16(11), 115011.

Froyd, C. A., Coffey, E. E., van der Knaap, W. O., van Leeuwen, J. F., Tye, A., & Willis, K. J. (2014). The ecological consequences of megafaunal loss: Giant tortoises and wetland biodiversity. Ecology Letters, 17(2), 144–154.

Gibbs, J. P., & Goldspiel, H. (2021). Population biology. In Galapagos giant tortoises (pp. 241–260). Elsevier.

Gibbs, J. P., Sterling, E. J., & Zabala, F. J. (2010). Giant tortoises as ecological engineers: A long-term quasi-experiment in the Galápagos Islands. Biotropica, 42(2), 208–214. 10.1111/j.1744-7429.2009.00552.x

Gounand, I., Harvey, E., Little, C. J., & Altermatt, F. (2018). Meta-ecosystems 2.0: Rooting the theory into the field. Trends in Ecology & Evolution, 33(1), 36–46.

Guézou, A., Trueman, M., Buddenhagen, C. E., Chamorro, S., Guerrero, A. M., Pozo, P., & Atkinson, R. (2010). An extensive alien plant inventory from the inhabited areas of Galapagos. PLoS One, 5(4), e10276.

Hartig, F. (2026). DHARMa: Residual Diagnostics for Hierarchical (Multi-Level / Mixed) Regression Models. 10.32614/CRAN.package.DHARMa

Holm-Hansen, O., & Riemann, B. (1978). Chlorophyll a determination: Improvements in methodology. Oikos, 438–447.

Holtgrieve, G. W., & Schindler, D. E. (2011). Marine-derived nutrients, bioturbation, and ecosystem metabolism: Reconsidering the role of salmon in streams. Ecology, 92(2), 373–385.

Hunter, E. A., Blake, S., Cayot, L. J., & Gibbs, J. P. (2021). Role in ecosystems. In Galapagos giant tortoises (pp. 299–315). Elsevier.

Jones, C. G., Lawton, J. H., & Shachak, M. (1994). Organisms as Ecosystem Engineers. Oikos, 69(3), 373–386. 10.2307/3545850

Kays, R., Davidson, S. C., Berger, M., Bohrer, G., Fiedler, W., Flack, A., Hirt, J., Hahn, C., Gauggel, D., Russell, B., Kölzsch, A., Lohr, A., Partecke, J., Quetting, M., Safi, K., Scharf, A., Schneider, G., Lang, I., Schaeuffelhut, F.,… Wikelski, M. (2022). The Movebank system for studying global animal movement and demography. Methods in Ecology and Evolution, 13(2), 419–431.

Keany, J. M., Burns, P., Abraham, A. J., Jantz, P., Makaga, L., Saatchi, S., Maisels, F., Abernethy, K., & Doughty, C. E. (2024). Using multiscale lidar to determine variation in canopy structure from African forest elephant trails. Remote Sensing in Ecology and Conservation, 10(5), 655–667.

Kitchell, J. F., O’Neill, R. V., Webb, D., Gallepp, G. W., Bartell, S. M., Koonce, J. F., & Ausmus, B. S. (1979). Consumer regulation of nutrient cycling. BioScience, 29(1), 28–34.

Kranstauber, B., Smolla, M., & Scharf, A. K. (2026). move: Visualizing and Analyzing Animal Track Data. 10.32614/CRAN.package.move

Leroux, S. J., & Loreau, M. (2008). Subsidy hypothesis and strength of trophic cascades across ecosystems. Ecology Letters, 11(11), 1147–1156.

Lipps, W. C., Braun-Howland, E. B., & Baxter, T. E. (Eds.). (2023). Standard Methods for the Examination of Water and Wastewater (24th ed.). APHA Press.

Loreau, M., Mouquet, N., & Holt, R. D. (2003). Meta-ecosystems: A theoretical framework for a spatial ecosystem ecology. Ecology Letters, 6(8), 673–679.

Lovich, J. E., Ennen, J. R., Agha, M., & Gibbons, J. W. (2018). Where Have All the Turtles Gone, and Why Does It Matter? BioScience, 68, 771–781.

Lundgren, E. J., Ramp, D., Stromberg, J. C., Wu, J., Nieto, N. C., Sluk, M., Moeller, K. T., & Wallach, A. D. (2021). Equids engineer desert water availability. Science, 372(6541), 491–495. 10.1126/science.abd6775

Marleau, J. N., Guichard, F., & Loreau, M. (2014). Meta-ecosystem dynamics and functioning on finite spatial networks. Proceedings of the Royal Society B: Biological Sciences, 281(1777), 20132094.

McInturf, A. G. et al. (2019). Vectors with autonomy: What distinguishes animal-mediated nutrient transport from abiotic vectors? Biological Reviews, 94(5), 1761–1773.

Metsio Sienne, J., Buchwald, R., & Wittemyer, G. (2014). Differentiation in mineral constituents in elephant selected versus unselected water and soil resources at Central African bais (forest clearings). European Journal of Wildlife Research, 60(2), 377–382.

Meysman, F. J., Middelburg, J. J., & Heip, C. H. (2006). Bioturbation: A fresh look at Darwin’s last idea. Trends in Ecology & Evolution, 21(12), 688–695.

Moore, J. W. (2006). Animal ecosystem engineers in streams. BioScience, 56(3), 237–246.

Naiman, R. J., Bilby, R. E., Schindler, D. E., & Helfield, J. M. (2002). Pacific salmon, nutrients, and the dynamics of freshwater and riparian ecosystems. Ecosystems, 5(4), 399–417.

Naiman, R. J., & Rogers, K. H. (1997). Large animals and system-level characteristics in river corridors. BioScience, 47(8), 521–529.

Pike, K. N., Blake, S., Gordon, I. J., Cabrera, F., Rivas-Torres, G., Laso, F. J., & others. (2023). Navigating agricultural landscapes: Responses of critically endangered giant tortoises to farmland vegetation and infrastructure. Landscape Ecology, 38, 501–516.

Polis, G. A., Anderson, W. B., & Holt, R. D. (1997). Toward an Integration of Landscape and Food Web Ecology: The Dynamics of Spatially Subsidized Food Webs. Annual Review of Ecology, Evolution, and Systematics, 28, 289–316. 10.1146/annurev.ecolsys.28.1.289

Pringle, R. M., Abraham, J. O., Anderson, T. M., Coverdale, T. C., Davies, A. B., Dutton, C. L., Gaylard, A., Goheen, J. R., Holdo, R. M., Hutchinson, M. C., & others. (2023). Impacts of large herbivores on terrestrial ecosystems. Current Biology, 33(11), R584–R610.

R Core Team. (2024). R: A Language and Environment for Statistical Computing. R Foundation for Statistical Computing. https://www.R-project.org/

Schmitz, O. J., Wilmers, C. C., Leroux, S. J., Doughty, C. E., Atwood, T. B., Galetti, M., Davies, A. B., & Goetz, S. J. (2018). Animals and the zoogeochemistry of the carbon cycle. Science, 362(6419), eaar3213.

Somaweera, R., Nifong, J., Rosenblatt, A., Brien, M. L., Combrink, X., Elsey, R. M., Grigg, G., Magnusson, W. E., Mazzotti, F. J., Pearcy, A., & others. (2020). The ecological importance of crocodylians: Towards evidence-based justification for their conservation. Biological Reviews, 95(4), 936–959.

Somveille, M., & Ellis-Soto, D. (2022). Linking animal migration and ecosystem processes: Data-driven simulation of propagule dispersal by migratory herbivores. Ecology and Evolution, 12, e9383.

Subalusky, A. L., Dutton, C. L., Njoroge, L., Rosi, E. J., & Post, D. M. (2018). Organic matter and nutrient inputs from large wildlife influence ecosystem function in the Mara River, Africa. Ecology, 99(11), 2558–2574.

Subalusky, A. L., Dutton, C. L., Rosi-Marshall, E. J., & Post, D. M. (2015). The hippopotamus conveyor belt: Vectors of carbon and nutrients from terrestrial grasslands to aquatic systems in sub-Saharan Africa. Freshwater Biology, 60(3), 512–525.

Subalusky, A. L., & Post, D. M. (2019). Context dependency of animal resource subsidies. Biological Reviews, 94(2), 517–538.

Tapia Aguilera, W., & Gibbs, J. P. (2023). Rewilding giant tortoises engineers plant communities at local to landscape scales. Conservation Letters, 16, e12968.

Tucker, M. A., Böhning-Gaese, K., Fagan, W. F., Fryxell, J. M., Van Moorter, B., Alberts, S. C., Ali, A. H., Allen, A. M., Attias, N., Avgar, T., Bartlam-Brooks, H., Bayarbaatar, B., Belant, J. L., Bertassoni, A., Beyer, D., Bidner, L., van Beest, F. M., Blake, S., Blaum, N.,… Mueller, T. (2018). Moving in the Anthropocene: Global reductions in terrestrial mammalian movements. Science, 359(6374), 466–469. 10.1126/science.aam9712

Vanni, M. J. (2002). Nutrient cycling by animals in freshwater ecosystems. Annual Review of Ecology and Systematics, 33(1), 341–370.

Wilcove, D. S., & Wikelski, M. (2008). Going, Going, Gone: Is Animal Migration Disappearing? PLoS Biology, 6(7), e188.

Yackulic, C. B., Blake, S., & Bastille-Rousseau, G. (2017). Benefits of the destinations, not costs of the journeys, shape partial migration patterns. Journal of Animal Ecology, 86(4), 972–982. 10.1111/1365-2656.12679

